# Phagocytosis is differentially regulated by LPS in M1- and M2-like macrophages via PGE_2_ formation and EP4 signaling

**DOI:** 10.1101/2024.09.29.615656

**Authors:** Rebecca Kirchhoff, Michel André Chromik, Nils Helge Schebb

## Abstract

Phagocytosis is a key process in human innate immune response. Human macrophages are important phagocytes engulfing and neutralizing pathogens and cell debris. In addition, they modulate the inflammatory process by releasing cytokines and lipid mediators. However, the link between oxylipins and phagocytosis in different macrophage phenotypes remains poorly understood.

In order to better understand the link between phagocytosis and the arachidonic acid (ARA) cascade, we established a phagocytosis assay in primary human ‘inflammatory’ M1- and ‘anti-inflammatory’ M2-like macrophages from PBMCs, representing extremes of macrophage phenotypes. The branches of the ARA cascade were investigated by quantitative targeted proteomics and metabolomics.

M1-like macrophages show a higher abundance of cyclooxygenase (COX)-2 and its products particularly after LPS stimulus compared to M2-like macrophages. LPS increased phagocytosis in M2-like, but not in M1-like macrophages. We demonstrate that the COX product PGE_2_ modulates the differential effects of LPS on phagocytosis: Via the EP4 receptor PGE_2_ signaling suppresses phagocytosis in primary human macrophages. Thus, blockage of COX, e.g. by NSAID, leads to an increase of phagocytosis also in ‘inflammatory’ M1-like macrophages and may shift the macrophages towards a more pro-resolving phenotype. This supports the well-described anti-inflammatory effects of these drugs.

## Introduction

Macrophages are important cells of the first line innate immunity and involved in host defense. Contact to pathogens such as gram-negative bacteria initiates the inflammatory response leading to the release of cytokines, chemokines (1), and activation of cytosolic phospholipase A_2_ (2). Cytosolic phospholipase A_2_ catalyzes the liberation of polyunsaturated fatty acids (PUFA) from membrane phospholipids (3). These PUFA can be converted via enzymatic and non-enzymatic pathways resulting in a large number of different oxylipins such as prostanoids, hydroperoxy-, mono and multi hydroxy-PUFA as well as epoxy-PUFA (4). Several of those are key regulators of the inflammatory process: Arachidonic acid (ARA)-derived prostaglandins (PG) formed by cyclooxygenase (COX) activity and leukotrienes formed by 5-lipoxygenase (LOX) activity are well-investigated pro-inflammatory mediators (4, 5) involved in pain, fever and asthma (6). However, several multi hydroxy-PUFA are discussed to have anti-inflammatory effects involved in the active resolution of inflammation (7) but remain controversial (8).

In addition to the release of signaling molecules, macrophages are capable of directly neutralizing pathogens by phagocytosis. This process is not only important for killing bacteria, but also contributes to the active resolution of inflammation: By ingesting apoptotic cells and cell debris, they prevent secondary necrosis and thus enable healing and restoration of healthy tissue (9). Hence, macrophages participate in both, the initiation and subsequent resolution of inflammation. Due to their high cellular plasticity, they adapt their functions in response to microenvironmental stimuli (10). This leads to different phenotypes such as inflammatory (M1) and anti-inflammatory (M2) macrophages (11). However, it is widely accepted that macrophages represent a spectrum of activated phenotypes, rather than discrete stable subpopulations (12). Differences in the phenotypes involve abundance of the enzymes of the ARA cascade and result in distinct oxylipin patterns (13, 14). However, the ARA cascade enzymes and oxylipins contribute to the regulation of phagocytosis in different macrophage phenotypes remains poorly understood.

Here, we investigated how the inflammatory stimulus bacterial lipopolysaccharide (LPS) impacts phagocytosis in primary human ‘inflammatory’ M1- and ‘anti-inflammatory’ M2-like macrophages, representing extremes of macrophage phenotypes. Particularly the role of the COX branch of the ARA cascade in the regulation of phagocytosis was evaluated. PGE_2_ signaling via the EP4 was found to be a key regulator suppressing the increase of phagocytosis in inflammatory M1-like macrophages.

## Materials and methods

### Chemicals and biological material

Human AB plasma was obtained from the blood donation center at University Hospital Düsseldorf (Düsseldorf, Germany). Lymphocyte separation medium 1077 was bought from PromoCell (Heidelberg, Germany). Recombinant human GM-CSF, M-CSF, IFNγ and IL-4 produced in *Escherichia coli* were purchased from PeproTech Germany (Hamburg, Germany). RPMI 1640 cell culture medium, L-glutamine and penicillin/streptomycin (5.000 units penicillin and 5 mg/mL streptomycin), LPS from *E. coli* (0111:B4), dextran 500 from *Leuconostoc spp.*, Cytochalasin D from *Zyosporium masonii* and copper sulfate pentahydrate were purchased at Merck (Taufkirchen, Germany). Latrunculin A, prostaglandin E_2_ (PGE_2_), PF-04418948 and GW627368 were bought from Cayman Chemical (local distributor Biomol, Hamburg, Germany). Accutase, DMSO and formaldehyde were obtained from Carl Roth (Karlsruhe, Germany). Resazurin was purchased from SERVA Electrophoresis GmbH (Heidelberg, Germany). Fluorescence-labeled polystyrene microspheres (latex beads) were bought as 2% or 2.5% solutions from Merck (Taufkirchen, Germany) and Thermo Fisher Scientific (Langenselbold, Germany). BCA reagent A was obtained from Fisher Scientific (Schwerte, Germany). The ultra-pure water with a conductivity of >18 MΩ*cm was generated by the Barnstead Genpure Pro system from Thermo Fisher Scientific (Langenselbold, Germany). Heavy labeled (lys: uniformly labeled (U)-^13^C_6_; U-^15^N_2_; arg: U-^13^C_6_; U-^15^N_4_) peptide standards were purchased from JPT Peptides (Berlin, Germany).

### Isolation and differentiation of primary human macrophages

Primary human macrophages were isolated as described (15). In brief, PBMC were isolated from buffy coats provided by blood donations at the University Hospital Düsseldorf, Germany or at Deutsches Rotes Kreuz West, Hagen, Germany. Blood was drawn with the informed consent of healthy human subjects and the study was approved by the Ethical Committee of the University of Wuppertal. PBMC were isolated by dextran (5%) sedimentation for 30-45 min and subsequent centrifugation (800 × *g* without deceleration, 10 min, 20 °C) on lymphocyte separation medium. The leucocyte ring was isolated, washed twice with PBS and cells were seeded in 60.1 mm² dishes in RPMI medium (1% P/S, 1% L-glutamine) in a humidified incubator at 37 °C and 5% CO_2_ for 1 h. Non-adherend cells were removed by washing, and RPMI medium (1% P/S, 1% L-glutamine) supplemented with 5% human AB plasma was added to the cells. For polarization towards M1-like macrophages, 10 ng/mL granulocyte-macrophage colony-stimulating factor (GM-CSF) was added to the medium for 7 days and additional 10 ng/mL interferon γ (IFNγ) for the final 48 h. For polarization towards M2-like macrophages, 10 ng/mL macrophage (M)-CSF was added for 7 days and additional 10 ng/mL interleukin 4 (IL-4) for the final 48 h.

### Phagocytosis assay

A schematic overview over the phagocytosis assay workflow is shown in **Fig. 1**. After differentiation, cells were harvested using Accutase (incubation for 20 min at 37 °C). Cells were transferred into 96-well plates at 50,000 cells/well using 100 µL cell culture medium. After resting overnight, cells were preincubated with test substances or vehicle control (0.1% DMSO) for 2-4 h at 37 °C. Cytotoxic effects of the tested substances were excluded by resazurin (almar blue) assay (16) and lactate dehydrogenase assay **(Fig. S1)** and only non-toxic concentrations were used. For phagocytosis, fluorescence-labeled polystyrene microspheres (beads) **(Tab. S1)** were used. Only beads having a fluorescence readout proportional to their concentration were used. After washing, the beads were opsonized in human plasma for 1.5-2 h at 37 °C.

**Fig. 1:**
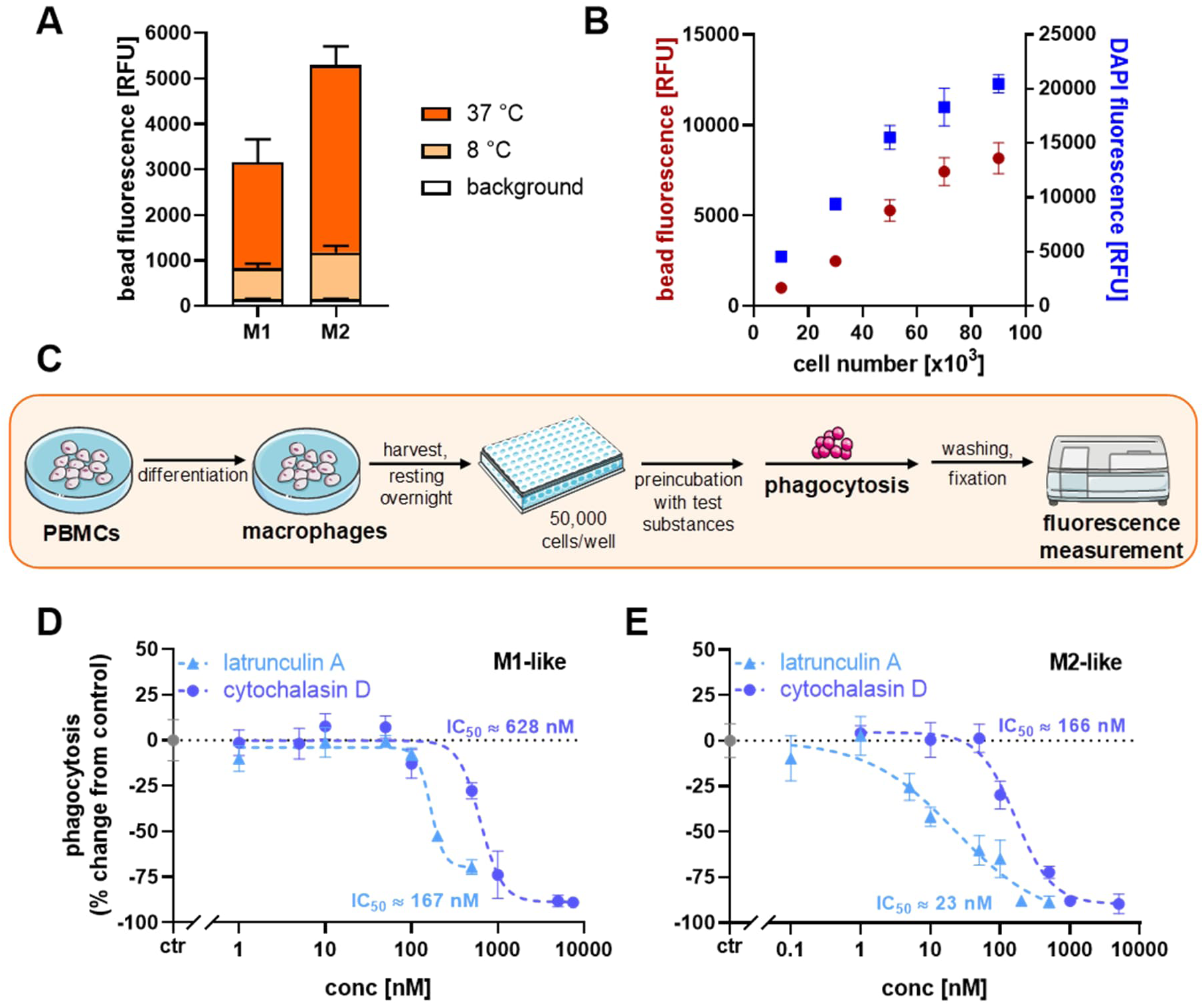
**Characterization of phagocytosis in M1- and M2-like macrophages**. Human monocytes were differentiated into M1-like or M2-like macrophages using 10 ng/mL GM-CSF or M-CSF (M1-/M2-like) for 7 days and additional 10 ng/mL IFNγ or IL-4 (M1-/M2-like) for the last 2 days. (**A**) Phagocytosis was carried out at 8 °C or 37 °C using 50,000 cells per well for 2 h. Shown is bead fluorescence at 610 nm after excitation at 575 nm as mean ± SD for n = 6-9 technical replicates from a pool of 3 subjects. (**B**) Phagocytosis in M2-like macrophages using different cell numbers, for 2 h followed by DAPI staining. Shown is the bead and DAPI fluorescence for the different cell numbers as mean ± SD for n = 8 replicates from a pool of 3 subjects. (**C**) Schematic overview over phagocytosis assay. (**D-E**) Dose dependent inhibition of phagocytosis in M1-like and M2-like macrophages by Latrunculin A or Cytochalasin D. Substances were preincubated at the indicated concentrations for 2 h. Shown is phagocytosis as % change from control (grey) as mean ± SD for n = 5-6 replicates from a pool of 4 subjects. The inhibitory potencies (IC_50_) were calculated based on %phagocytosis relative to vehicle control (0.1% DMSO).

For the phagocytosis assay, supernatants of the 96-well plate were discarded, and opsonized bead solution, which had been diluted with medium (6.25 ·10^-3^% beads, 5% human plasma), was added together with the preincubated test substance to the cells for 2 h at 37 °C in a humidified incubator. Phagocytosis was stopped by discarding the supernatants and carefully washing the cells three times with 100 µL PBS. Cells were fixated using 4% formaldehyde for 10 min at 37 °C. After three times washing with 100 µL PBS, fluorescence of beads was measured using a plate reader (Infinite 200 PRO, Tecan). For DAPI staining, 10 µg/mL DAPI was added to the cells for 15 min at room temperature followed by three times washing with PBS and fluorescence measurement with the plate reader (λ_ex_ 358 nm, λ_em_ 461 nm).

### Quantification of oxylipin and protein levels by LC-MS/MS

Differentiated macrophages were harvested from cell culture dishes via the cold shock method (15) and dry pellets were stored at -80 °C until use. Oxylipin and protein levels were analyzed from the same cell pellet. Cell pellets were resuspended in MeOH/H_2_O (50/50, *v*/*v*) containing antioxidant solution (0.2 mg/mL BHT, 100 μM indomethacin, 100 μM soluble epoxide hydrolase inhibitor trans-4-[4-(3-adamantan-1-yl-ureido)-cyclohexyloxy]-benzoic acid (t-AUCB) in MeOH) (17, 18), sonicated and protein content was determined via bicinchoninic acid assay (BCA) (19). Internal standards (IS) for oxylipin analysis were added to the samples before proteins were precipitated using MeOH for at least 30 min at -80 °C, followed by centrifugation (20 000 × *g*, 10 min, 4 °C). The supernatant was used for the oxylipin analysis and the precipitated protein pellet for the analysis of the protein level (13).

Oxylipin determination was carried out according previously published methods (17, 18). In brief, oxylipins were purified by solid phase extraction (SPE) on a non-polar (C8)/strong anion exchange mixed mode material (Bond Elut Certify II, 200 mg, Agilent, Waldbronn, Germany), and analyzed using an 1290 Infinity II LC system (Agilent), coupled to a QTRAP mass spectrometer operated in electrospray ionization (ESI(-)) mode (Sciex, Darmstadt, Germany) operated in scheduled selected reaction monitoring. Quantification was carried out by external calibration using IS (13, 20).

Quantitative protein analysis was carried out by LC-MS/MS as described (13, 14, 21). In brief, the protein pellet was dissolved in 5% (*w*/*v*) sodium deoxycholate with protease inhibitor (100/1, *v*/*v*), proteins were precipitated using acetone and centrifuged (15 000 × *g*, 10 min, 4 °C). After subsequent incubations with 200 mM dithiothreitol, 200 mM iodoacetamide, and again 200 mM dithiothreitol, 10 µg/mL trypsin (trypsin-to-protein ratio of 1:50) was added to digest the proteins for 15 h. The digestion was stopped using acetic acid (pH 3-4). IS (heavy labeled peptides) were added and samples were purified on Strata-X SPE 33 µm Polymeric Reversed Phase cartridges (100 mg/3 mL, Phenomenex LTD, Aschaffenburg, Germany). Peptides were reconstituted in 50 µL 15% ACN/0.1% acetic acid and analyzed on an 1290 Infinity II LC system (Agilent) coupled to QTRAP instrument (Sciex) in ESI(+)-mode operated in scheduled selection reaction monitoring mode. Quantification was carried out by external calibration using IS (13, 21).

Analyst (Sciex, version 1.7) was used for instrument control and Multiquant software (Sciex, version 3.0.2) was used for data analysis. The concentrations of oxylipins and proteins were calculated based on the protein content determined via bicinchoninic acid assay (19).

Data are presented as mean ± standard error of mean (SEM) or standard deviation (SD) as indicated. Z-factor (z) was calculated using the means (µ) and standard deviations (σ) of the inhibitor latrunculin A (µ_neg_, σ_neg_) and of the LPS stimulation (µ_pos_, σ_pos_).

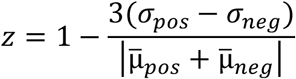

## Results

For the investigation of phagocytosis in human macrophages, we established and optimized a phagocytosis assay in primary human M1- and M2-like macrophages. Both types of macrophages showed phagocytic activity using fluorescence-labeled polystyrene microspheres (beads) **(Fig. 1A)**. In order to optimize the phagocytosis assay, several parameters of the assay, such as cell number, duration of phagocytosis, and number of wash steps, were carefully selected **(Fig. 1B, C, Fig. S2)**. Both, latrunculin A and cytochalasin D inhibited phagocytosis in M1- and M2-like macrophages in a dose-dependent manner, with IC_50_ values in the low nM range **(Fig. 1D, E)**. Overall, optimization resulted in a z factor higher than 0.5 indicating a good bioassay **(Fig. S3)**.

The inflammatory stimulus LPS was found to increase phagocytosis in M2-like macrophages in a dose- and time-dependent manner **(Fig. 2A, B)**. Incubation times between 2 to 24 h and concentrations between 10-100 ng/mL resulted in the highest increase in phagocytosis (1.6-fold). In contrast, no effect of LPS on phagocytosis was observed in M1-like macrophages **(Fig. 2C)**. In order to elucidate the reasons for these differences, ARA enzyme abundance in the two types of macrophages after LPS stimulation was analyzed and revealed 1.6-fold higher COX-2 abundance in the M1-like type compared to M2-like **(Fig. 3A)**. In line with this, concentrations of COX products were approximately 2-fold higher **(Fig. 3B)**. Therefore, we hypothesized that the different effect of LPS on phagocytosis in the two macrophage types may be mediated by COX activity. In order to test this, the impact of COX inhibitors and dexamethasone on phagocytosis was evaluated **(Fig. 2C, D)**: Indeed, LPS increased phagocytosis in both macrophage types (M1-like: 1.4-fold, M2-like 1.7-fold) in the presence of the three compounds. Moreover, PGE_2_, one of the main COX products, inhibited phagocytosis in both macrophage types in a dose-dependent manner (IC_50_(M1-like) ≈ 2.5 nM, IC_50_(M2-like) ≈ 17 nM) **(Fig. 4A, B)**. Among the four PGE_2_ receptors, EP2 and EP4 are present in the macrophages (22–24). Using an antagonist against EP4, inhibition of phagocytosis by PGE_2_ was restored, whereas the antagonist against EP2 showed no effect **(Fig. 4C)**. This demonstrates that phagocytosis is regulated via EP4 signaling in human M1- and M2-like macrophages.

**Fig. 2:**
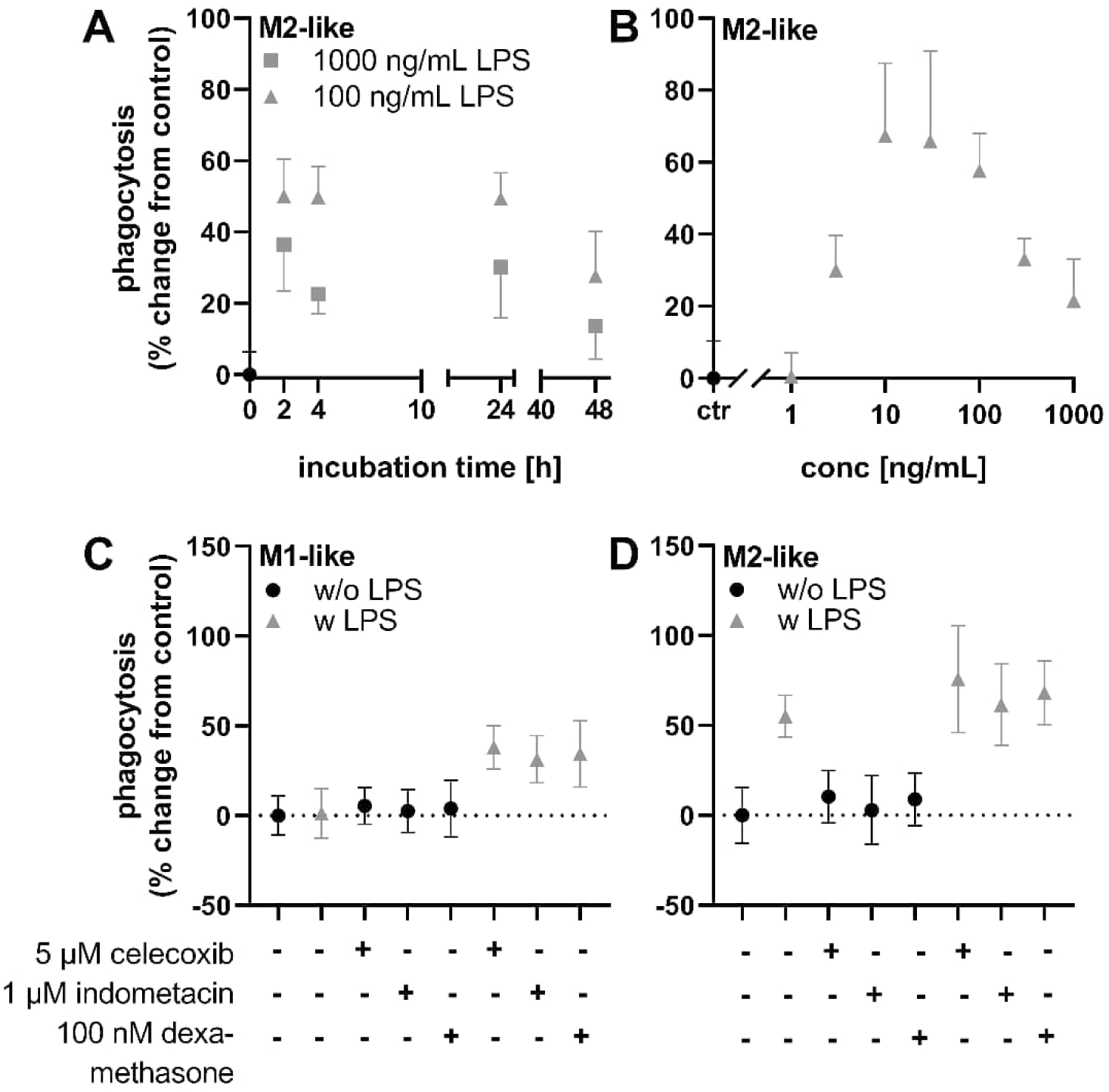
Phagocytosis is differentially modulated by LPS in M1- and M2-like macrophages. **(A)** M2-like macrophages were preincubated with 0.1 or 1 µg/mL LPS for different periods of time or **(B)** the indicated concentrations for 4 h. **(C)** M1-like or **(D)** M2-like macrophages were preincubated with Celecoxib (5 µM), Indometacin (1 µM) or Dexamethasone (100 nM) for 30 min before addition of 0.1 µg/mL LPS for 4 h. Shown is phagocytosis as % change from vehicle control (0.1% DMSO) as mean ± SD for *n* = 5-6 replicates from a pool of 3-4 subjects.

**Fig. 3:**
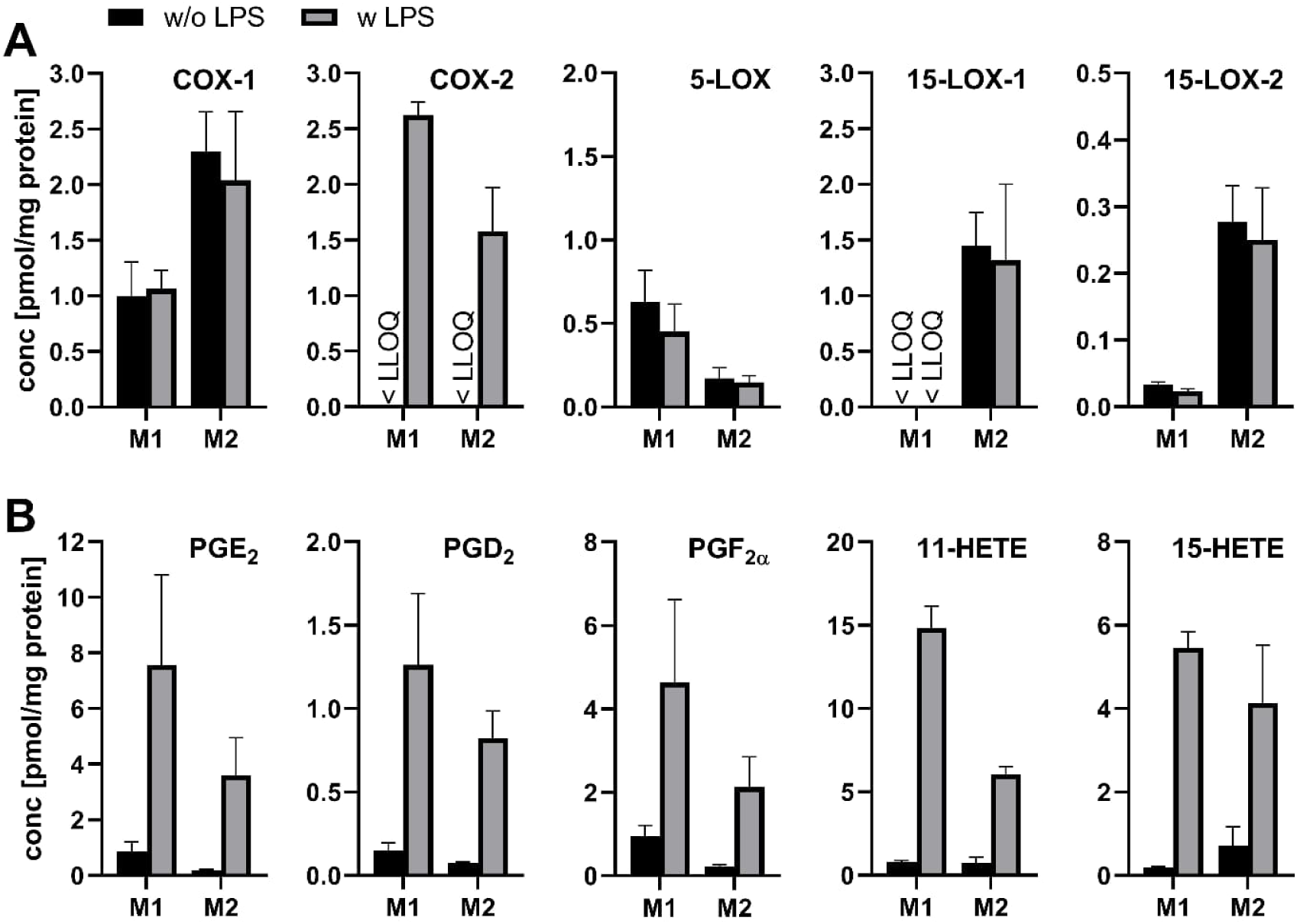
ARA enzyme abundance and COX activity of macrophages with and without LPS stimulation. **(A)** Macrophages were analyzed for ARA enzyme abundance by quantitative proteomics and **(B)** free oxylipins by quantitative metabolomics with (w) and without (w/o) additional LPS (0.1 µg/mL) treatment for 6 h. Shown are mean ± SEM for *n* = 3 subjects.

**Fig. 4:**
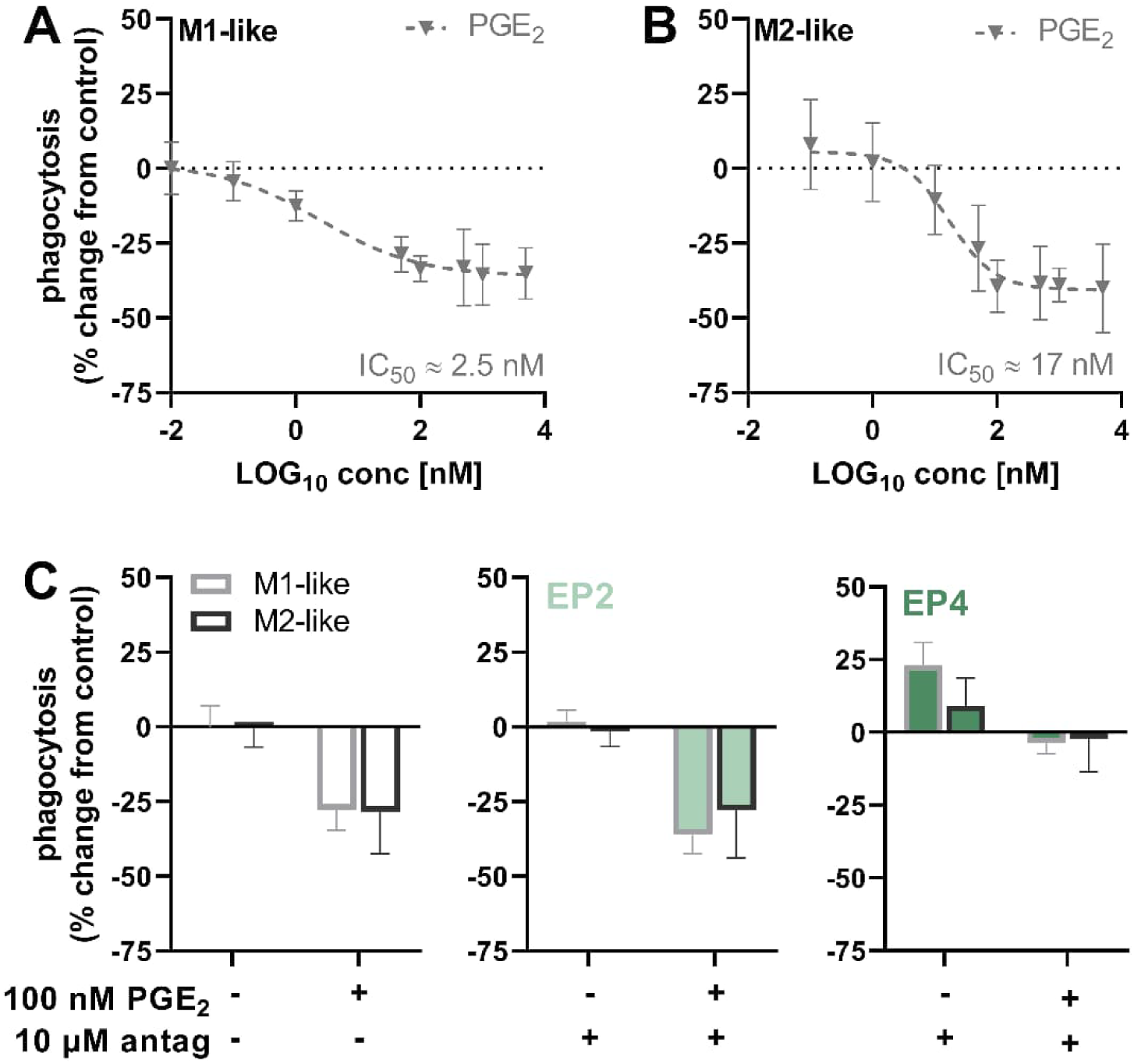
Regulation of phagocytosis by COX-product PGE_2_. **(A)** M1-like or **(B)** M2-like macrophages were incubated with different PGE_2_ concentrations for 2 h. **(C)** Macrophages were treated with 100 nM PGE_2_ for 2 h or EP2 antagonist (PF-04418948, 10 µM) or EP4 antagonist (GW627368, 10 µM). Antagonists were tested alone and in combination with 100 nM PGE_2_ and added 30 min before PGE_2_ incubation. Shown is phagocytosis as % change from vehicle control (0.1% DMSO) as mean ± SD for *n* = 5-6 replicates from a pool of 3-4 subjects.

## Discussion

Phagocytosis is a key process of innate immune cells such as macrophages to eliminate foreign invaders and cell debris during inflammatory processes. Here, we show the optimization of a phagocytosis assay to investigate, how the COX branch of the ARA cascade is involved in the regulation of phagocytosis.

### Optimization of the phagocytosis assay

For the investigation of phagocytosis, a variety of methods have been described using different bacteria/particles to be engulfed. The readout includes labor- and cost-intensive techniques such as confocal microscopy and flow cytometry (25, 26). Moreover, simple readouts based on fluorescence measurements using a plate reader and a 96-well plate format are described (27–29). Here, we established such an assay with fluorescence-labeled polystyrene microspheres (beads) for primary human macrophages. Using the amount of nucleic acids stained with DAPI, the cell number was examined indirectly and optimized for the assay **(Fig. 1B)** resulting in a sufficient high cell number leading to an optimal fluorescence readout, but ensuring sufficient space for the cells. Further parameters such as resting time after transfer of cells to 96-well plates, number of cells per well, the length of phagocytosis, and number of required washing steps to remove > 90% of not internalized beads **(Fig. S2)**, were optimized. However, analysis of the fluorescence signals after phagocytosis at 8 °C **(Fig. 1A)** showed that beads cannot completely be removed by washing. This is consistent to other reports showing that beads are not engulfed, but adhere to the cell surface at 8 °C (27). Therefore, we calculate fluorescence/phagocytosis data relative to a vehicle control and established controls for the assay.

Comparing the differently polarized macrophage types, M2-like macrophages showed a 1.5-fold higher phagocytic activity than M1-like macrophages **(Fig. 1A)**. This is in line with previous reports (30, 31) and could be due to the high abundance of scavenging receptors in M2-like macrophages which support phagocytic functions (32, 33).

Cytochalasin D and latrunculin A, well-known inhibitors of phagocytosis were able to inhibit phagocytosis in a dose-dependent manner, whereby maximum inhibition was achieved at lower concentrations for latrunculin A (200 nM) than for cytochalasin D (1 µM) **(Fig. 1D, E)**. M2-like macrophages were more sensitive to the inhibitors as indicated by lower IC_50_ values **(Fig. 1D, E)**. Similar inhibitory concentrations (12-120 nM) were reported for latrunculin A for mouse peritoneal macrophages (34) and higher concentrations (10-40 µM) for cytochalasin D for mouse peritoneal macrophages and for J774A.1 cells, a murine macrophage cell line (35, 36) which may be explained by the different cell type. Cytochalasin D and latrunculin A are suitable chemical probes used as control to inhibit phagocytosis in the assay.

### Phagocytosis is differentially modulated by LPS in M1- and M2-like macrophages

Information on substances that reliably enhance phagocytosis is limited in the literature, in contrast to the well-described inhibitors. We investigated LPS, since it has been widely described, that LPS impacts a variety of macrophage functions such as phagocytosis (37–39). Indeed, in M2-like macrophages, LPS increased phagocytosis in a dose- and time-dependent manner **(Fig. 2A, B)**. In contrast, LPS showed no effect on phagocytosis in M1-like macrophages **(Fig. 2C)**. Interestingly, both – stimulation and inhibition of phagocytosis – have been described in the literature following a stimulation with LPS (39–43): Whereas LPS increased phagocytosis in mouse bone-marrow derived macrophages and human monocytes using similar conditions (1 h or 20-24 h, using concentrations of 10-100 ng/mL) (40, 43), phagocytosis was suppressed in mouse peritoneal macrophages (39, 41). This leaves the role of LPS in the regulation of phagocytosis controversial. In order to elucidate the differential modulation of phagocytosis by LPS in the two macrophage types, the involvement of ARA cascade enzymes was investigated.

The ARA cascade enzyme pattern showing higher levels of COX-1, 15-LOX-1 and -2 and lower levels of 5-LOX in M2-like compared to M1-like macrophages **(Fig. 3B)** and was comparable to previous reports (13, 14). LPS stimulation led to elevated *PTGS2* expression in both macrophage types, leading to a 1.6-fold higher COX-2 abundance in the M1-like type **(Fig. 3A)**. The COX activity was relevantly higher in M1-like macrophages as well as the levels of its products such as PGE_2_, PGD_2_ and PGF_2α_ were 2-4-fold higher in M1-like macrophages at basal levels and after stimulation with LPS (e.g., PGE_2_ M1 vs. M2, basal: 0.85 ± 0.35 vs 0.20 ± 0.02 pmol/mg protein; LPS: 7.57 ± 3.25 vs 3.59 ± 1.36 pmol/mg protein) **(Fig. 3B)**. This was in line with levels of 11-HETE **(Fig. 3B)**, which is a well-known side product of COX-2 activity (44, 45). 15-HETE showed basal higher concentrations in M2-like macrophages compared to M1-like presumably due to 15-LOX-1 activity; after LPS stimulus levels were increased. This can be explained by COX-2 activity, with 15-HETE being one of the main side products (44, 45). Both, enzyme abundance and oxylipin levels underline that M2-like macrophages have lower COX-2 abundance and activity after LPS stimulus compared to M1-like macrophages. Therefore, we hypothesized that COX (products) may contribute to the differential effect of LPS on phagocytosis observed in M1-like macrophages (no effect) compared to M2-like macrophages (stimulation).

In order to investigate the impact of COX activity on phagocytosis, COX-inhibitors and dexamethasone were tested: The tested substances alone had no effect on phagocytosis **(Fig. 2C, D)** which fits to the previous finding that basal COX activity is very low in both macrophage types. However, in combination with LPS, phagocytosis was increased in both macrophage types (M1-like, 35%; M2-like, 69%). Similar findings, an upregulation of phagocytosis, were already reported for murine bone marrow derived macrophages and alveolar macrophages treated with celecoxib (46, 47), for rat alveolar macrophages treated with celecoxib or indomethacin (48, 49), as well as for human monocytes with dexamethasone (50). This indicates that (some) COX products counteract the stimulating effect of LPS. Thus, the strong upregulation of COX-2 abundance and activity by LPS in M1-like macrophages can explain why LPS could not increase phagocytosis in M1-like macrophages. However, besides induction of COX-2, LPS induces a variety of other changes in the cells that are discussed affecting phagocytosis such as cytokines (40), number of specific surface receptors (39) and the cytoskeletal network (39). Further research is needed to elucidate which other mechanisms of action of LPS are involved in the regulation of phagocytosis in M1- and M2-like macrophages.

PGE_2_ was the highest abundant prostaglandin in M1-like macrophages after LPS stimulus **(Fig. 3B)**. Thus, we tested if the LPS effect is mediated by this prostaglandin. PGE_2_ inhibited phagocytosis in a dose-dependent manner in both macrophage types with a half-maximal inhibition (IC_50_) in the low nM range **(Fig. 4A, B)**. Observed IC_50_ values were slightly lower for M1-like indicating these cells were more sensitive to PGE_2_. Our results are in line with the majority of the studies which reported an inhibitory effect of PGE_2_ on phagocytosis *in vitro* (40, 48, 49, 51–55) and *in vivo* (56, 57). Moreover, the extent of inhibition of phagocytosis by PGE_2_ up to 40% is comparable to values found for other monocyte/macrophage systems (approximately 25-85%) shown in the literature (40, 49, 53, 55).

PGE_2_ affects changes in cell functions through activation of distinct G protein-coupled E prostanoid (EP) receptors (EP1-EP4). Among the EP receptors, EP2 and EP4 are present in human macrophages (22–24). In order to determine, which of the two receptors is involved in the regulation of phagocytosis by PGE_2_ in human M1- and M2-like macrophages, we tested specific antagonists against EP2 and EP4 **(Fig. 4C)**. The EP2 antagonist PF-04418948 did not affect phagocytosis by PGE_2_ in both cell types. In contrast, EP4 antagonist GW627368 entirely blocked the inhibitory effect of PGE_2_. This indicates, that exogenously added PGE_2_ mediates its inhibitory effects on phagocytosis through EP4 activation in human macrophages. GW627368 alone increased phagocytosis in M1-like macrophages (by approximately 23%), suggesting that already basal formed levels of PGE_2_ (without LPS stimulus) are high enough to elicit inhibitory effects on phagocytosis. This fits to the comparatively high levels of approximately 1 pmol/mg protein PGE_2_ in M1-like macrophages determined by LC-MS/MS **(Fig. 3A)**. However, the unselective COX inhibitor indomethacin alone did not increase phagocytosis in M1-like macrophages, indicating it was not able to completely block PGE_2_ formation or residual levels of PGE_2_ were present prior to inhibitor treatment.

Both receptors, EP2 and EP4, signal predominantly via stimulatory G protein (G_s_), activating adenylyl cyclase (AC) and increasing the intracellular levels of cAMP (58). There is evidence that PGE_2_ inhibits phagocytosis via a cAMP dependent pathway (49, 51, 54, 55, 59), which fits to our findings. However, reports are inconsistent, which of the EP receptors are involved in signal transduction: For rat alveolar macrophages and mouse bone marrow derived macrophages, an EP2 dependent signaling was reported (47, 49, 51). However, other studies reported the involvement of both receptors in rat alveolar macrophages (48) and in THP-1 derived macrophages (54) indicating that the PGE_2_ signaling may be highly cell type dependent.

Our results indicate, that PGE_2_ signaling on phagocytosis is mainly through EP4 receptor in primary human macrophages. The different PGE_2_ levels in M1- and M2-like macrophages as a result of different COX-2 activities are involved in the modulation of phagocytosis by LPS. This shows the importance of the link between the COX branch of the ARA cascade and regulation of phagocytosis in human macrophages and provides a mechanistical explanation of the well-investigated anti-inflammatory effects of an COX-2 inhibition (60): By suppressing the formation of pro-inflammatory PGE_2_, e.g., by using COX inhibitors, phagocytosis can be increased which in turn limits inflammation, enables tissue restoration and may shift the M1-like macrophages towards a more pro-resolving phenotype.

## Acknowledgment

This work was supported by the German Research Foundation (SCHE 1801) to NHS and a Ph.D. fellowship from the Friedrich-Ebert-Stiftung to RK.

## Ethics approval

Peripheral blood monocytic cells (PBMC) were isolated from buffy coats obtained from blood donations. Blood samples were drawn with the informed consent of the human subjects. The study was approved by the Ethical Committee of the University of Wuppertal.

## Source of biological material

PBMC were isolated from buffy coats obtained from blood donations at the University Hospital Düsseldorf and Deutsches Rote Kreuz-blood donation service West (Hagen).

## Abbreviations

ARA: arachidonic acid
COX: cyclooxygenase
EP: prostaglandin E_2_ receptor
ESI: electrospray ionization
FA: fatty acid
HETE: hydroxy eicosatetraenoic acid
IS: internal standard
LOX: lipoxygenase
MS: mass spectrometry
P/S: penicillin/streptomycin
PUFA: polyunsaturated fatty acid

## Supporting Information

**Table S1:**
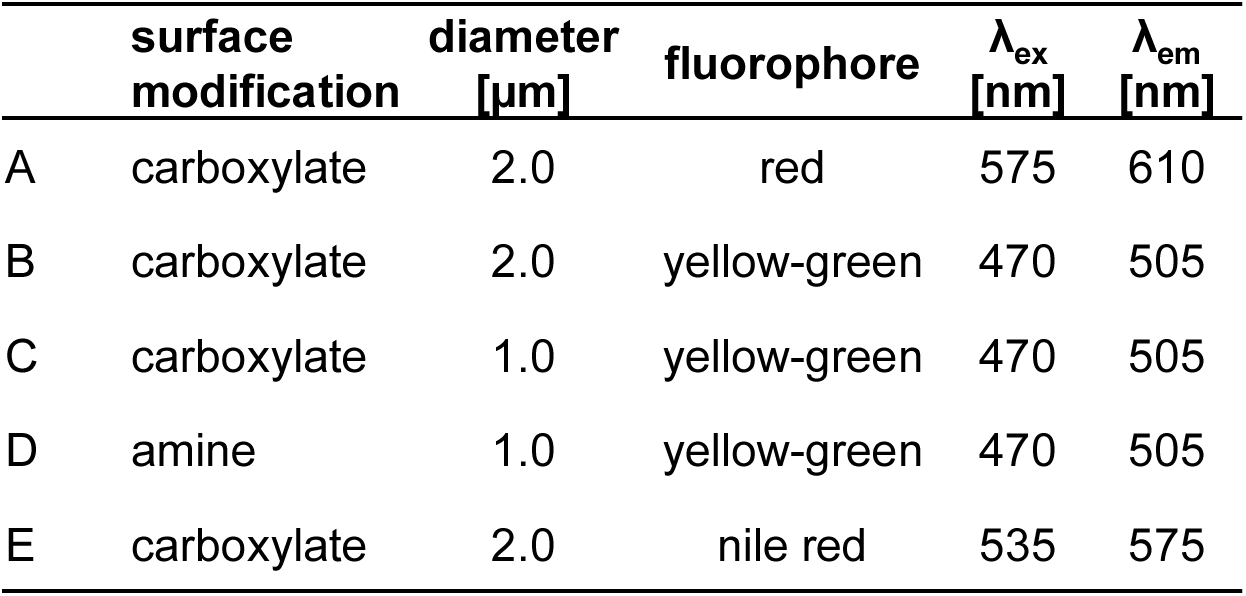
Properties of the fluorescent labeled polystyrene microspheres (beads) tested in the phagocytosis assay.

| | surface<br>modification | diameter<br>[$\mu\text{m}$] | fluorophore | $\lambda_{\text{ex}}$<br>[nm] | $\lambda_{\text{em}}$<br>[nm] |
| --- | --- | --- | --- | --- | --- |
| A | carboxylate | 2.0 | red | 575 | 610 |
| B | carboxylate | 2.0 | yellow-green | 470 | 505 |
| C | carboxylate | 1.0 | yellow-green | 470 | 505 |
| D | amine | 1.0 | yellow-green | 470 | 505 |
| E | carboxylate | 2.0 | nile red | 535 | 575 |

**Figure S1:**
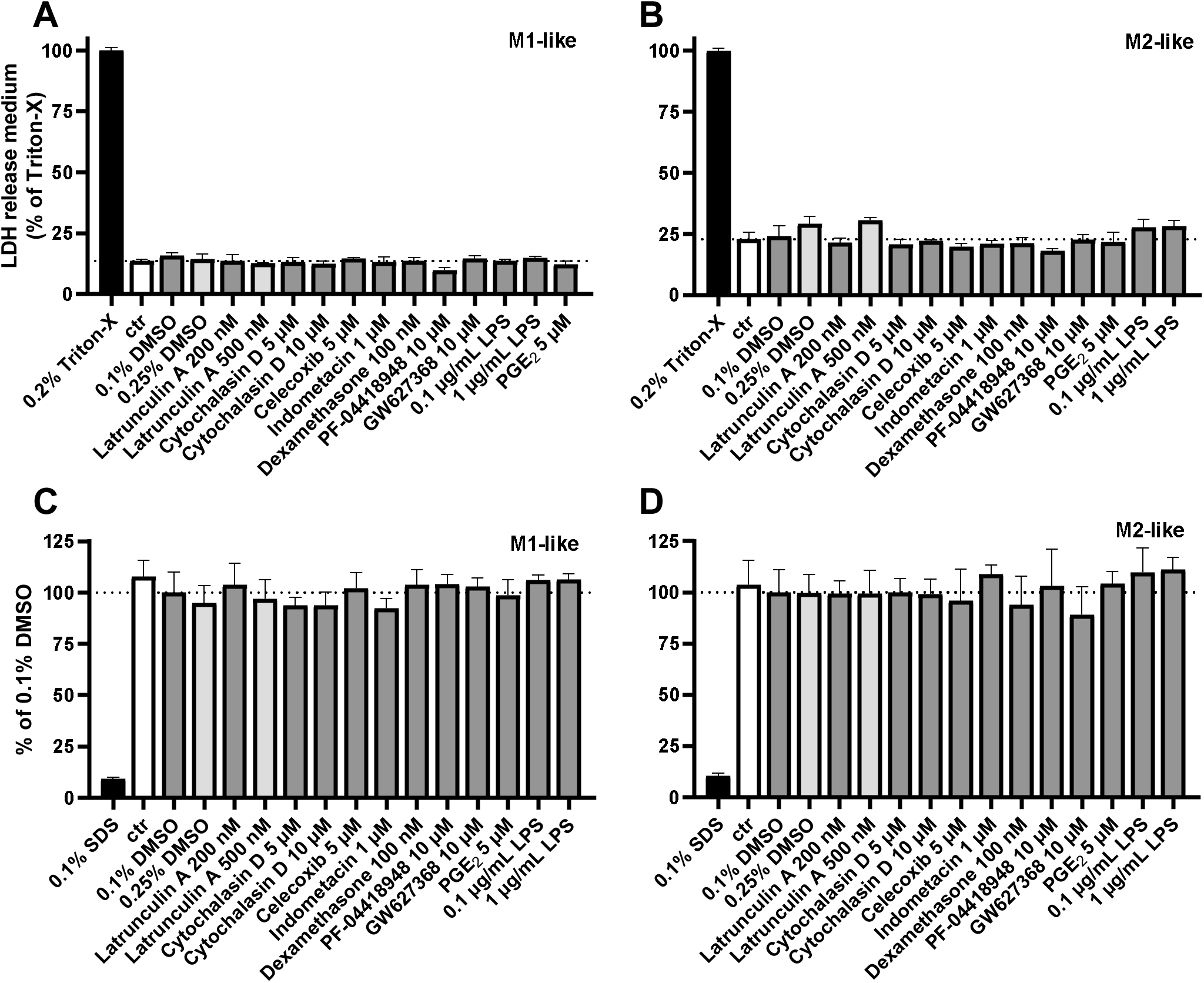
**Effect of the tested compounds on cell viability determined by lactate dehydrogenase (LDH) leakage (A, B) or metabolic activity by resazurin assay (C, D) in primary human macrophages**. Human monocytes were differentiated into M1-like or M2-like macrophages using 10 ng/mL GM-CSF or M-CSF (M1-/M2-like) for 7 days and additional 10 ng/mL IFNγ or IL-4 (M1-/M2-like) for the last 2 days. The cells were incubated with compounds at the indicated concentrations for 4 h. DMSO served as vehicle control and Triton X (LDH assay) or SDS (resazurin assay) as positive control. Dehydrogenase activity in the medium was measured by the decrease in absorbance at 340 nM for 45 min during the NADH dependent reduction of pyruvate to lactate. LDH leakage was determined by comparing LDH activities in culture medium and lysed cells. Shown are mean ± SD for n = 3 replicates from a pool of 3 subjects. Dehydrogenase activity was measured as resorufin formation by fluorometric readout at 590 nm after excitation at 530 nm. Shown are mean ± SD for *n* = 3-6 replicates from a pool of 3 subjects.

**Figure S2:**
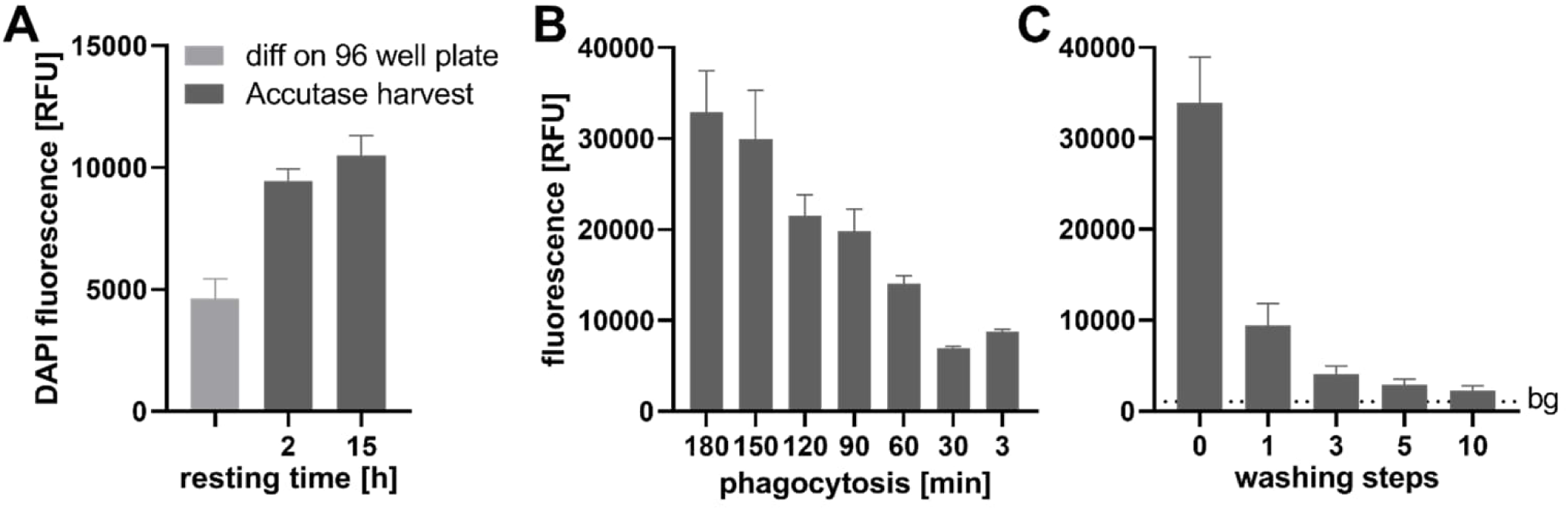
Characterization of resting time of cells prior phagocytosis assay, duration of phagocytosis assay and number of washing steps. Primary human monocytes were differentiated into M2-like macrophages using 10 ng/mL M-CSF for 7 days and additional 10 ng/mL IL-4 for the last 2 days. **(A)** Cells were either differentiated directly in 96 well plates (light grey) or cultured and differentiated on dishes followed by harvest using Accutase (dark grey) and transferred to 96 well plates (50,000 cells/well). The cells were allowed to rest/adhere to the plates for 2 or 15 h. The colonization efficacy (cell number) was determined based on the amount of DNA determined by incubation with DAPI. The cells were washed with PBS and fluorescence (λ_ex_ 358 nm, λ_em_ 461 nm) was measured in order to determine the optimal duration of the phagocytosis. **(B)** Cells were incubated with beads for different periods of time ranging between 3 to 180 min. After washing with PBS, bead fluorescence was analyzed in the cells. **(C)** After phagocytosis assay (for 120 min) cells were washed different times with PBS as indicated and bead fluorescence was measured in the cells in order to determine the necessary number of washing steps to remove adherent, non-ingested beads. All results are shown as mean ± SD, *n* = 5-9 technical replicates from a pool of 3 human subjects.

**Figure S3:**
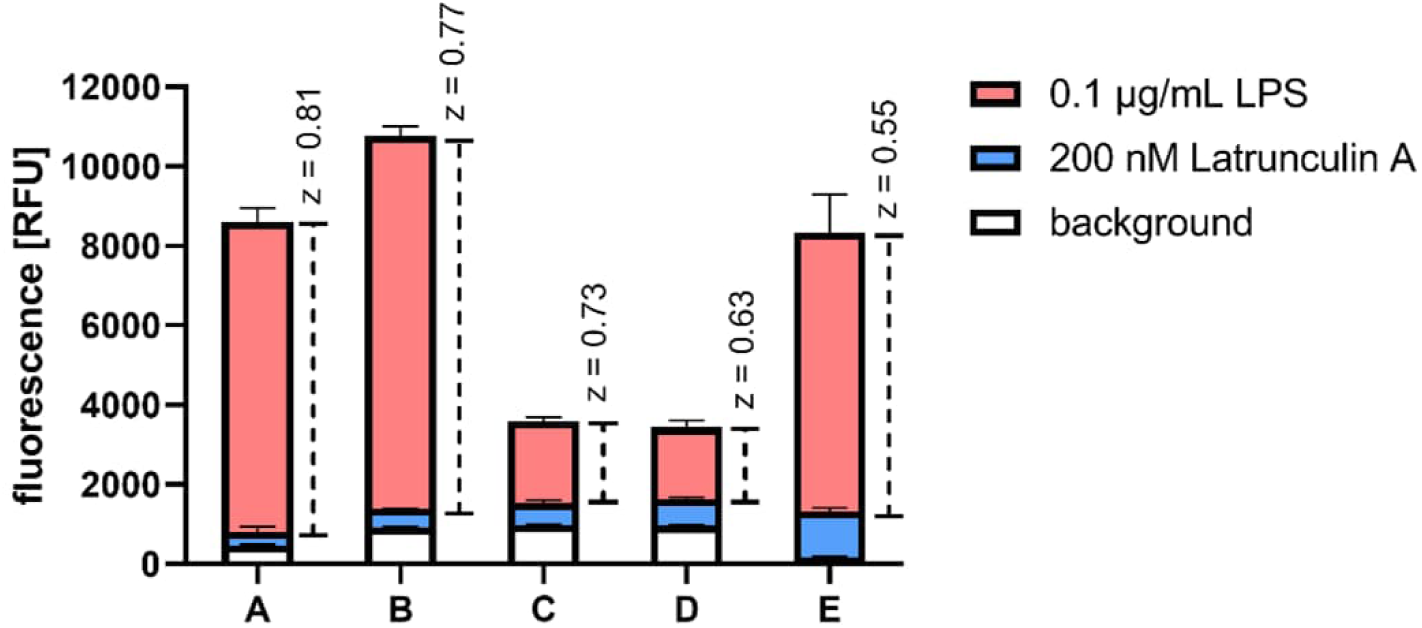
Impact of type of fluorescent beads on the performance of the phagocytosis assay in M2-like macrophages. Primary human monocytes were differentiated into M2-like macrophages using 10 ng/mL M-CSF for 7 days and additional 10 ng/mL IL-4 for the last 2 days. Cells were preincubated with 0.1 µg/mL LPS (positive control) for 4 h or 200 nM Latrunculin A (negative control) for 2 h before starting phagocytosis. Phagocytosis was carried out using different beads A-D (Tab. S1) for 2 h. Results are shown as mean ± SD, *n* = 5-9 technical replicates from a pool of 3 human subjects. Z-Factors were calculated based on mean and standard deviation of signals of positive control and negative control.

